# The specificity of fatigue cerebral substrates in early Multiple Sclerosis - An fMRI study

**DOI:** 10.1101/2025.06.27.661966

**Authors:** Camille Guillemin, Maëlle Charonitis, Nikita Beliy, Florence Requier, Gaël Delrue, Emilie Lommers, Gilles Vandewalle, Pierre Maquet, Christophe Phillips, Fabienne Collette

**Author notes:** Correspondance: Fabienne Collette, GIGA-CRC in vivo Imaging Université de Liège, Allée du 6 Août n°8 4000 Liège, Belgium.

## Abstract

**Context:** The cerebral substrates of fatigue in patients with Multiple Sclerosis (pwMS) are not elucidated yet. This study aims at exploring the disease-specific functional brain substrates of fatigue in pwMS with a recent disease history

**Methods:** Sixteen pwMS (disease history < 5 years) and 17 matched Healthy Controls (HC) performed a N-Back task with three difficulty levels during fMRI acquisitions following high vs. low fatigue induction. Measures of subjective trait and state fatigue were also recorded. Behavioral performance at n-back task and evolution of subjective fatigue states were analyzed by means of Bayesian repeated measures analyses of variance. Functional MRI data were analyzed to determine between-group differences (1) in task-related brain activity, independently of trait fatigue score; (2) in the association between trait fatigue and brain activity

**Results:** A similar trajectory was observed in the two groups for subjective and task-related measures following fatigue induction. No between-group difference was observed in brain activity unrelated to fatigue score. However, negative associations between trait fatigue score and brain activity were observed in pwMS, while the associations are positive in HC. Specifically, interactions between group and task difficulty were observed in regions belonging to the Striato-Thalamo-Cortical (STC) network and the fronto-parietal cortex

**Conclusion:** Group-specific patterns of brain activity related to fatigue were identified in pwMS and HC in the STC circuitry, despite similar behavioral performance and subjective fatigue level. This confirms the implication of the STC loops in fatigue pathophysiology, occurring from the early stages of the disease.

## Introduction

Fatigue is one of the most frequent and debilitating symptoms of Multiple Sclerosis (MS; Bakshi et al., 2000; Weiland et al., 2015). Despite its high prevalence, the pathophysiology of MS fatigue is not completely established. Fatigue state is partly explained by contributing symptoms such as depression, anxiety and sleep disorders (Weiland et al., 2015). Lesion load and cortical atrophy are also linked to fatigue complaints in people with multiple sclerosis (pwMS), albeit not systematically (see ARM, Ribbons, Lechner-Scott, & Ramadan, 2019).

A central cause for MS fatigue is often put forward. In 1997, Roelcke and colleagues (1997) observed decreased glucose metabolism in frontal lobes, premotor cortices, right supplemental motor area and basal ganglia, as well as an increased metabolism in cingulate cortex and cerebellum, which were associated to fatigue level in pwMS. Accordingly, Chaudhuri and Behan (2000, 2004) proposed that fatigue results from a disruption of the basal ganglia functions. The basal ganglia are extensively connected to the cortex and thalamus (DeLong & Wichmann, 2010; Rǎ dulescu, Herron, Kennedy, & Scimemi, 2017) through Striato-Thalamo-Cortical loops (STC, sometimes referred as Cortico-Striato-Thalamo-Cortical loops) (Alexander, DeLong, & Strick, 1986; Peters, Dunlop, & Downar, 2016). If their exact number remains debated, three main STC circuits are classically identified: the motor loop, with inputs from the sensorimotor cortex, the associative (or cognitive) loop, with main inputs from the dorso-lateral prefrontal cortex, and the limbic (or affective/motivational) loop, involving the anterior cingulate cortex, medial orbitofrontal cortex, the hippocampus and amygdala (DeLong & Wichmann, 2010; Posner et al., 2014).

STC loops have been implicated in the pathophysiology of MS fatigue (ARM et al., 2019; Newland, Starkweather, & Sorenson, 2016; Palotai & Guttmann, 2020). Resting-state and motor fMRI studies identified both hyper-and hypo-activity patterns within STC networks. Fatigue related hypo-activity was interpreted as reflecting dysfunction of the basal ganglia and structural alterations along the cortico-cortical and cortico-subcortical circuitry, while hyperactivity in the STC and the default mode networks was generally interpreted as relating to a costly functional reorganization of brain networks leading to fatigue (ARM et al., 2019).

Oddly enough, fatigue was never explored by task-fMRI in recently diagnosed patients with MS. Yet, fatigue occurs at any disease stages, even following the first demyelinating episode (Runia, Jafari, Siepman, & Hintzen, 2015) despite low lesion load and disability. Since MS fatigue appears to be distinct from fatigue sensation experienced physiologically by healthy subjects (Krupp, Alvarez, LaRocca, & Scheinberg, 1988), an identification of its cerebral correlates at disease onset would be of great interest.

Here, we aim at exploring the functional substrates of fatigue in pwMS with a recent disease history. Following a fatigue induction protocol, we compared fatigue-independent and fatigue-dependent brain activity of pwMS and matched healthy controls (HC) during a cognitive task with three levels of difficulty.

## Methods

### Participants

Sixteen recently diagnosed pwMS and 17 HC matched for sex, age and education were included (see the detailed protocol on OSF for a complete chart flow of participants recruited: https://osf.io/egr6d/?view_only=cdcc343c7d71406685b038e46e88145b).

Patients presented either a Relapsing-Remitting (RR) or a Clinically Isolated Syndrome (CIS) course of the disease, according to the 2017 McDonald criteria (Thompson et al., 2018). Inclusion criteria for pwMS were absence of relapse in the last 6 months, disease duration below or equal to 5 years and a score at the Expanded Disability Status Scale (EDSS, Kurtzke, 1983) under 4. Exclusion criteria for both groups comprised the existence of other neurological or psychiatric diseases, a history of mild or severe traumatic brain injury, the use of medication affecting fatigue state and/or alertness, substance abuse, color blindness, native language other than French and age above 45 years old. The following medications were considered as an exclusion criterion if they were not stopped at least 5 half-lives prior participation to the study: benzodiazepine, neuroleptic, tricyclic antidepressant, beta blocker, anticholinergic and antiepileptic drugs.

### General procedure

This study was approved by the local ethics committees of the University Hospital of Liège (B707201835630). Participants received detailed information on the protocol and gave written informed consent prior to their participation to this study. Participants were asked to observe a stable sleep-wake cycle during their participation to the study, with a sleep duration of at least 7 hours per night and to avoid stimulating beverages 24 hours prior to an appointment. To limit circadian confounds, appointments for a single participant were scheduled at the same time of the day, according to her/his preferences.

At visit 1, participants filled-in questionnaires and were administered a training and calibration of the Time Load Dual Back task, which is a dual task (TLDB, Borragán, Slama, Bartolomei, & Peigneux, 2017). During sessions 2 and 3, cognitive fatigue was induced with the TLDB task performed in two conditions, counterbalanced across sessions: the High or Low Cognitive Load conditions (HCL and LCL, respectively). Following fatigue induction, fMRI data were acquired in order to assess the effect of cognitive load (i.e. during fatigue induction) on brain activity while performing a cognitive task of varying difficulty (the N-Back task, Kirchner, 1958).

### Behavioral data

#### Questionnaires

Several questionnaires were administered, assessing mood status (Beck Depression Inventory (BDI-II, Beck, Ward, Mendelson, Mock, & Erbaugh, 1961), anxiety level (State-Trait anxiety inventory: STAI, Spielberger, Gorsuch, Ushene, Vagg, & Jacobs, 1983), sleep quality (Pittsburgh Sleep Quality Index: PSQI, Buysse, Reynolds, Monk, Berman, & Kupfer, 1998), daytime sleepiness (Epworth Sleepiness Scale: ESS, Johns, 1991) and fatigue (including the Fatigue Scale for Motor and Cognitive Functions: FSMC, Penner et al., 2009).

Score at the FSMC is one of our primary measures of interest. The FSMC is a 20-items scale with a great reliability and validity designed to assess fatigue symptom in MS population in a holistic way (Penner et al., 2009). This scale also provides sub-scores for motor and cognitive fatigue, as well as cut-off scores for normal, mild, moderate and severe fatigue. We decided to use the trait fatigue score (i.e. every-day fatigue) from FSMC rather than state fatigue (i.e. fatigue sensation at the moment) as measured by a visual analogue scale since FSMC captures a more complete and consistent measure of fatigue symptom across subjects. As scores at FSMC subscales are highly correlated, the total score was selected for fMRI analysis.

#### Fatigue induction protocol: the Time Load Dual Back (TLDB) task

The TLDB is a computerized dual task combining a number parity judgment task and a classical N-Back working memory task. Cognitive load induced by the task can be manipulated easily while maintaining the same task complexity by adjusting the time available to process stimuli (i.e. the Stimulus Time Duration, STD). STD was individually calibrated before the experimental condition and corresponds to the fastest presentation rate at which accuracy remained above 85%. By doing so, the STD was tailored to the subject’s processing abilities. This point is particularly relevant here, as pwMS frequently present deficits in information processing speed and working memory (Chiaravalloti & DeLuca, 2008).

For each participant, the TLDB was presented for 32 minutes in two conditions during two separate and counterbalanced sessions: the High Cognitive Load session (HCL) and the Low Cognitive Load session (LCL). In the HCL condition, stimuli were presented at a pace corresponding to the STD, while in the LCL condition the pace was slowed down by 50%. Results from the TLDB have been reported elsewhere (see Guillemin et al., 2022).

#### The N-Back task

Following fatigue induction, participants underwent an fMRI session to determine brain activity during a computerized N-Back task (Kirchner, 1958) with three levels of difficulty (1 to 3-Back, See Supplemental Figure S1). In the N-Back task, letters are presented and participants had to detect if the current letter matched the one presented N-position(s) before (with a yes/no response). The 1-Back condition is generally considered as a reference level allowing control for brain activity associated to perceptual, motor and short-term memory processes. The task was composed of 18 successive blocks of 16 stimuli (6 blocks per difficulty level), presented in pseudorandom order. Stimuli were displayed in large white font on a black background. Each block started with instructions displayed during 5000ms. In each block, letters were presented with a time presentation of 1700ms followed by an inter-stimulus fixation cross of 500ms. At the end of the block, a fixation cross of 12000ms was displayed before the beginning of the next block. The total task duration was of 16 minutes, approximately. Participants were trained outside and inside the scanner until they felt comfortable with the task and key-responses.

#### Subjective state

Subjective state of the participants was assessed at 3 different time-points per session: before (T0) and after (T1) fatigue induction by TLDB task, and after the fMRI N-Back task (T2), including measurement of sleepiness, fatigue and motivation levels as assessed by the Karolinska Sleepiness Scale (KSS, Åkerstedt & Gillberg, 1990) and Visual Analogue Scales (VAS, Lee, Hicks, & Nino-Murcia, 1991).

#### Statistical analysis

Behavioral data analysis was conducted using Bayesian statistical inference (Keysers, Gazzola, & Wagenmakers, 2020) using the software JASP (Jeffreys’s Amazing Statistics Program v.0.16; https://jasp-stats.org). Results are reported in terms of Bayes Factor (BF) which corresponds to the likelihood ratio of evidence provided by the data over two hypotheses (BF_10_ represents the marginal likelihood of the alternative hypothesis H1 over the one of the null hypothesis H0. BFi_ncl_ represents the likelihood of adding an effect in the statistical model over the null model). Significant results were determined using Jeffreys’s grades of evidence (Jeffreys, 1961). In short BF_10/incl_ values <1 are in favor of the null hypothesis/model and values >1 in favor of the alternative hypothesis/tested model (see Table 1 for presentation of level of evidence with BF10 as example). Between groups differences in questionnaires were investigated using Mann-Whitney tests; performance at the N-back task as well as results to subjective scales (KSS & VAS) were analyzed by means of Analysis of Variance for repeated measures (rmANOVAs).

**Table 1.** Jeffreys’s Bayes factor evidence category (Jeffreys, 1961)

| BF<sub>10</sub> | Interpretation |
| --- | --- |
| >100**** | Extreme evidence for H1 |
| 30–100*** | Very Strong evidence for H1 |
| 10–30** | Strong evidence for H1 |
| 3–10* | Moderate evidence for H1 |
| 1–3 | Anecdotal evidence for H1 |
| 1 | No evidence |
| 0.333–1 | Anecdotal evidence for H0 |
| 0.1–0.333* | Moderate evidence for H0 |
| 0.033–0.1** | Strong evidence for H0 |
| 0.01–0.033*** | Very Strong evidence for H0 |
| <0.01**** | Extreme evidence for H0 |

Level of evidence for the alternative (H1) and the null (H0) hypothesis depending on Bayes Factor (BF<sub>10</sub> here). For instance, a BF<sub>10</sub> of 5 indicates that the data observed are 5 times more likely to happen under the alternative hypothesis rather than the null, corresponding to a moderate evidence for H1. The same logic applies to BF<sub>incl</sub>, with BF<sub>incl</sub> above 3 indicating evidence of an effect of the variable in the studied model, and BF<sub>inc</sub> under 0.333 indicating an absence of effect.

### MRI data

#### Data acquisition and MRI sequences

Functional MRI time series were acquired on a 3T scanner (Magnetom PRISMA, Siemens Medical Solution, Erlangen, Germany) with a 64-channel head coil. Multislice T2*-weighted functional images were acquired with the multi-band gradient-echo echo-planar imaging sequence (CMRR, University of Minnesota) using axial slice orientation and covering the whole brain/most of the brain (36 slices, multiband factor = 2, FoV = 216x216 mm², voxel size 3x3x3 mm³, 25% interslice gap, matrix size 72x72x36, TR = 1170 ms, TE = 30 ms, FA = 90°). The three initial volumes were discarded to avoid T1 saturation effects. Respiration and pulse signals were recorded during fMRI time series in order to derive physiological regressors and correct for physiological noise in the BOLD signal. Additional field map data were acquired, consisting on two magnitude and a single phase difference images to correct the whole fMRI series for distortion (40 slices, voxel size 3x3x3 mm³, TR = 634 ms, TE1 = 10 ms, TE2 = 12.46 ms, FA = 90°). For anatomical reference, a high-resolution T1-weighted image was acquired for each subject (T1-weighted 3D magnetization-prepared rapid gradient echo (MPRAGE) sequence, TR = 1900 ms, TE = 2.19 ms, inversion time (TI) = 900 ms, FoV = 256x240 mm², matrix size = 256x240x224, voxel size = 1x1x1 mm³, acceleration factor in phase-encoding direction R=2). An additional T2 FLAIR sequence was acquired and used for lesion segmentation in the pwMS group (TR = 5000 ms, TE = 5.16 ms, inversion time (TI) = 1800 ms, FoV = 256x240 mm², matrix size = 256x240x224, voxel size = 1x1x1 mm³, acceleration factor in phase-encoding direction R=2).

#### Data preprocessing

MRI data were organized according to BIDS format (Gorgolewski et al., 2016, BIDS version 1.2, https://bids.neuroimaging.io/). Spatial preprocessing of all images was carried out using SPM12 (Statistical Parametric Mapping, Wellcome Trust Centre for Neuroimaging, http://www.fil.ion.ucl.ac.uk/spm). Lesion maps in the pwMS group were derived using the Lesion Segmentation Toolbox (LST, Schmidt et al., 2012), with the FLAIR image, then manually corrected, and verified by a neurologist. Anatomical images of HC subjects were first corrected for intensity bias using light regularization and a 30 mm Full Width at Half Maximum (FWHM) cutoff. Corrected FLAIR images were coregistered to anatomical corrected T1w images. Those images were then used for multi-channel segmentation using an extended tissue probability map (with high sensitivity to subcortical grey-matter volumes) and normalized onto the MNI space with forward deformation fields. The same procedure was applied to images of pwMS except that a lesion mask was added in the coregistration step and segmentation was performed using the Unified Segmentation with Lesion (USwithLesion, Phillips & Pernet, 2018) toolbox for SPM. Quality of segmentation was assessed visually for all subjects. EPI field maps were used to calculate Volume Displacement Maps (VDM), for each session separately. All EPI volumes were then realigned (registration to the first volume and estimation of movement parameters) and unwarped using the corresponding VDM. The mean unwarped EPI image was then coregistered to the anatomical T1w image, along with the realigned and unwarped volumes of both sessions. Finally, all time series were normalized onto the MNI space using the deformation field of the T1w image and smoothed with an 8mm FWHM Gaussian kernel. Additional visual quality control was performed on all smoothed images.

#### Statistical analysis

Functional MRI data were analyzed using a two-steps procedure taking into account intra-individual variance, then inter-individual variance (Figure 1). For each subject, a general linear model was used to compute statistical maps of changes in regional brain responses across fatigue induction conditions (HCL and LCL) as well as task difficulty (1 to 3 Back conditions). Blocks corresponding to 1-Back, 2-Back and 3-Back working memory loads in each fatigue condition were modeled using boxcar functions and convolved with a canonical hemodynamic response function. Motion and physiological regressors were added in the model and considered as covariates of no interest (see Fig. 1A). A high-pass filtering of 256 seconds was implemented in the matrix design to remove low-frequency drifts from the time series. Individual contrast maps were computed for the main effect of fatigue induction condition and task difficulty (HCL vs. LCL, HCL 1-Back, HCL 2-Back, HCL 3-Back, HCL 2-Back vs. HCL 1-Back, HCL 3-Back vs. HCL 1-Back).

The resulting summary statistic images were then entered into a second level two samples T-test model. Score at the total FSMC scale for trait fatigue was mean-centered across all participants and added as group-specific continuous covariate of interest in the model. An inclusive grey-matter mask was computed from tissue probability maps (probability of voxel belonging to grey matter above 0.15) and used to remove voxels of non-interest (such as in ventricles) and improve readability of results. First, group analyses (pwMS vs HC) on trait fatigue-independent activity (not associated with FSMC score) were performed with F-tests (Fig. 1B). Second, statistical maps of brain activity associated with the FSMC fatigue score (in one group or both groups) were extracted to define regions potentially associated to fatigue (*p* < 0.001 uncorrected, Fig. 1C). That set of regions was further used for small-volumes correction in the group comparison of correlations (Fig. 1D). This analysis identified the associations between fatigue and brain activity that significantly differ between groups (F-Test, *p*_FWE_ < 0.05, cluster forming threshold *p* < 0.001 uncorrected). Brain labels were based on the Automated Anatomical Labeling atlas (AAL; Tzourio-Mazoyer et al., 2002) and directionality of significant correlations was assessed graphically (see supplemental Figure S2).

**Figure 1.**
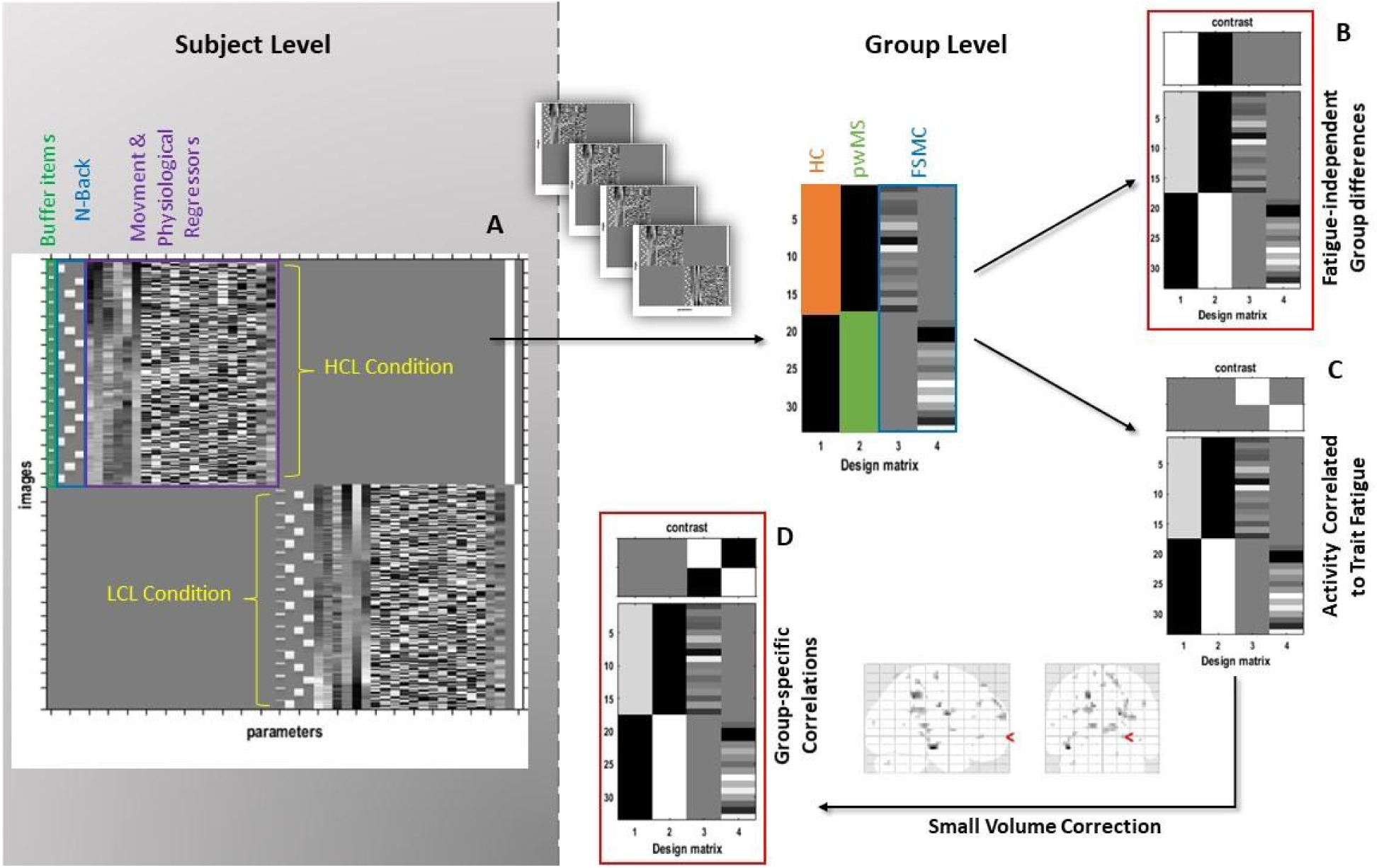
fMRI statistical analysis overview Contrast maps were first computed individually at the subject level. **(A)**, with a GLM including the two fMRI sessions (HCL and LCL fatigue induction conditions, yellow). 1B, 2B and 3B blocks (blue), as well as buffer items (green) were entered as individual regressors. Movement and physiological noise were added as regressors of no interest (purple). The resulting summary contrast maps were then entered into a F-Test group level analyses (middle of the Figure), with fatigue score added in covariable of interest (FSMC). Group differences **(B)** were assessed in trait fatigue-independent brain activity. Trait fatigue-dependent activity (**C – D**) was next assessed. First, statistical activity maps were computed in all subjects **(C**) and used as small volume correction for analysis **(D)** which explored if correlations with FSMC score were group-specific. Results from **(B)** and **(C)** are reported and discussed here (red, illustration of group comparison for the association between fatigue and brain activity).

## Data availability

Data from this study will be made available upon reasonable request from any qualified investigator. Supplementary analyses are planned with the dataset and a data transfer agreement will be signed before access to the requested data.

## Results

### Behavioral data

#### Demographics and questionnaires

No between-group difference (Table 2) was found regarding self-assessment questionnaires (all BF_10_ < 3), however BF_10_ did not allow to conclude regarding an absence of difference between groups, especially for the physical subscale of the FSMC and the score at the PSQI. Regarding the total score at the FSMC, 5 pwMS (31.25%) and 2 HC (11.77%) reached the cutoff score of 63 for severe fatigue, and 7 pwMS (43.75%) and 11 HC (64.71%) had mild to moderate fatigue.

**Table 2.** Demographic data and disease-related information.

|  | pwMS<br>(*n* = 16) | HC<br>(*n* = 17) | BF<sub>10</sub> |
| --- | --- | --- | --- |
| Age, y, mean(SD) | 31.44 (4.62) | 31.42 (5.99) | 0.338<sup>a</sup> |
| Women, *n* (%) | 12 (75%) | 13 (76.47%) | 0.356<sup>b</sup> |
| Education, y, mean (SD) | 14.19 (2.17) | 14.41 (1.46) | 0.385<sup>a</sup> |
| FSMC (median(SD), max = 100) | 57 (19.81) | 53 (14.97) | 0.381<sup>c</sup> |
| FSMC Motor (median(SD), max = 50) | 30.5 (10.29) | 26 (8.04) | 0.508<sup>c</sup> |
| FSMC Cognitive (median(SD), max = 50) | 28 (10.38) | 27 (10.38) | 0.353<sup>c</sup> |
| STAI – Y1 (median(SD), max = 80) | 33 (11.04) | 31 (8.45) | 0.347<sup>c</sup> |
| STAI – Y2 (median(SD), max = 80) | 43 (11.76) | 37 (12.94) | 0.381<sup>c</sup> |
| BDI-II (median(SD), max = 63) | 7 (12.02) | 8 (10.33) | 0.339<sup>c</sup> |
| PSQI (median(SD), max = 21) | 7.5 (2.73) | 5 (1.87) | 1.046<sup>c</sup> |
| ESS (median(SD), max = 24) | 10 (5.47) | 9 (4.32) | 0.362<sup>c</sup> |
| Time since first symptom, y, mean(SD) | 2.04 (0.95) | n.a. | / |
| Time since diagnosis, y, mean(SD) | 1.80 (1.22) | n.a. | / |
| EDSS, median(range) | 1.75 (1 – 2.5) | n.a. | / |

pwMS: person with multiple sclerosis; HC: Healthy controls; FSMC: Fatigue Scale for Motor and Cognitive function; STAI: State (Y1) Trait (Y2) Anxiety Inventory; BDI-II: Beck Depression Inventory; PSQI: Pittsburgh Sleep Quality Index; ESS: Epworth Sleepiness Scale. <sup>a</sup>independent sample T-test, <sup>b</sup>contingency table analysis for independent multinomial sampling (Jamil et al., 2017), <sup>c</sup>Mann-Whitney test.

#### Fatigue induction and subjective state

Bayesian ANOVAs showed a significant effect of time for all scales (VAS fatigue and motivation, KSS; Table 3). Post-hoc analyses (paired-sample T-tests with BF correction, Westfall, Johnson, & Utts, 1997) showed that fatigue and sleepiness increased after administration of the TLDB task (all corrected BF_10_ T0 vs T1 >7) and remained stable afterwards, until the end of the N-Back task (BF_10_ T1 vs T2 <0.2). On the contrary, motivation decreased after the TLDB task (corrected BF_10_ = 5143) and raised again after the N-back task, yet did not reach initial state (corrected BF_10_ T0 vs T2 = 3.388). A visual representation of the effects of time and fatigue induction on VAS and KSS is available in the supplemental material (Figure S3).

Importantly, fatigue induction condition (HCL vs LCL) did not influence subjective fatigue and sleepiness (BF_10_ < 0.33). Moreover, evolution of subjective state over time and across conditions followed a similar pattern in pwMS and HC (Condition*Time*Group interactions, all BF_10_ < 0.33).

**Table 3.** Effects of fatigue induction Condition, Time and Group on subjective states.

|  | BF<sub>inclusion</sub> |  |  |
| --- | --- | --- | --- |
|  | Fatigue | Sleepiness | Motivation |
| Condition | **0.169\*** | **0.169\*** | 0.352 |
| Time | **3.526\*** | **3984.73\*\*\*** | **1168.89\*\*\*** |
| Group | **0.149\*** | 0.353 | 0.375 |
| Condition*Time | 0.353 | **0.100\*\*** | **0.119\*** |
| Condition*Group | 0.449 | **0.245\*** | **0.216\*** |
| Time*Group | **0.121\*** | **0.295\*** | **0.283\*** |
| Condition*Time*Group | **0.256\*** | **0.183\*** | **0.214\*** |

Asterisks refer to the level of evidence as expressed in Table 1

#### N-Back task

Mean reaction times (RTs) to correct answers, hit rates (number of Hits divided by number of targets) and d’ sensitivity index were computed. Effects of Group, fatigue induction Condition and N-Back Difficulty level (1-to 3-Back) are reported in table 4. We observed an overall effect of task difficulty on all measures, with an increase of RTs and a decrease of hit rate and d’ index as difficulty increases (Figure 2). The fatigue induction Condition did not influence performance, with a moderate evidence for absence of effect regarding mean RTs and sensitivity. Evidence for all Group effects were inconclusive, with a substantial evidence for an absence of effect. Finally, most interaction effects show moderate evidence for an absence of effect. Overall, we can conclude that only task difficulty had an effect on performance in our protocol, with no clear evidence for an effect of fatigue induction condition nor disease.

**Figure 2.**
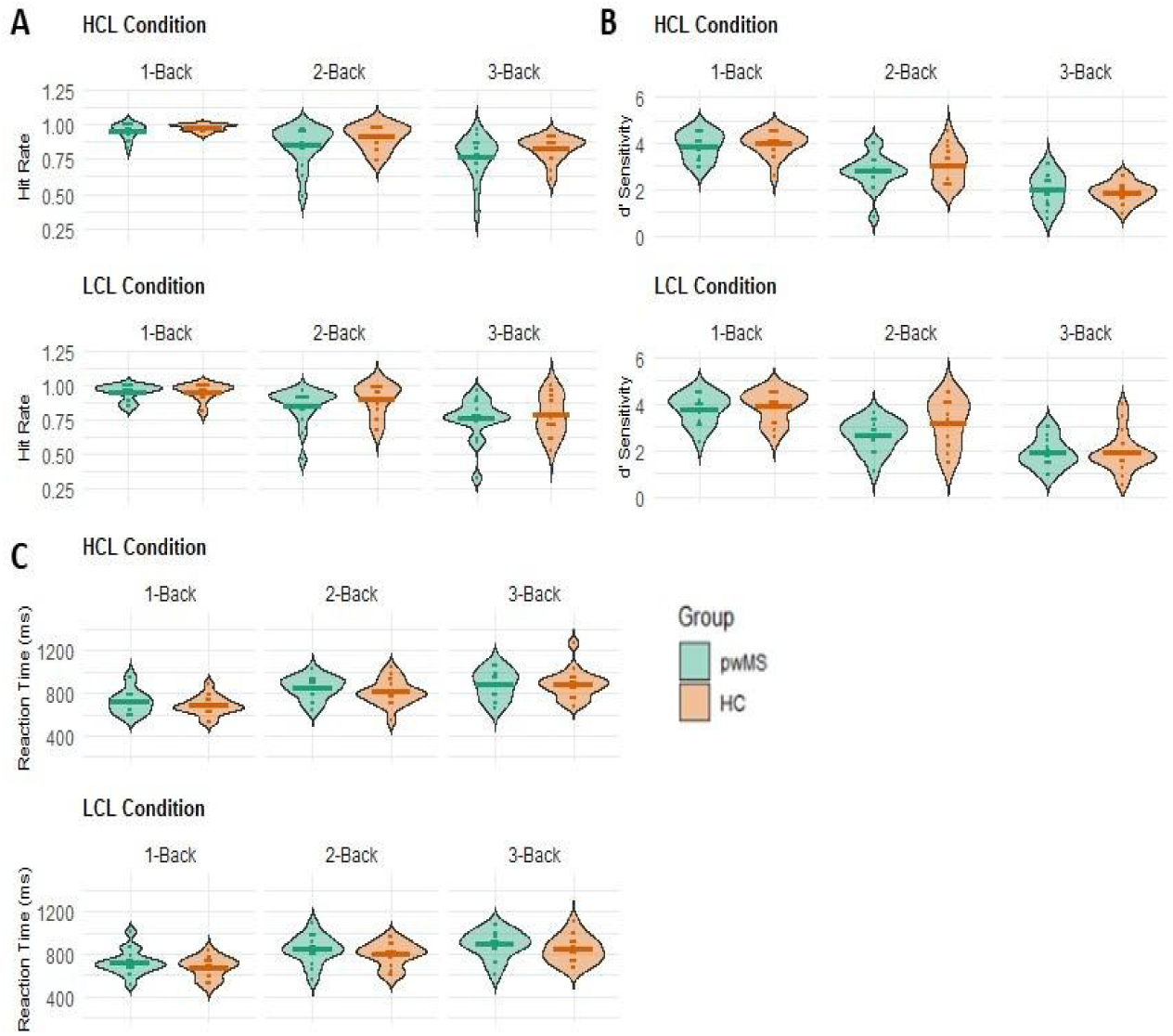
Performance at the fMRI N-Back task. Performance at the N-Back task for people with Multiple Sclerosis (pwMS, green violins) and Healthy Controls (HC, orange violins) expressed by Hit Rate (A), d’ sensitivity index (B) and mean Reaction Time (C) depending on task difficulty (1 to 3 Back) and fatigue induction (High Cognitive Load, HCL, upper panels and Low Cognitive Load, LCL, lower panels). Dots represent single subjects; horizontal bar represents group means.

**Table 4.** Effects of task Difficulty, fatigue induction Condition and Group on N-Back performance.

|  | BF<sub>inclusion</sub> |  |  |
| --- | --- | --- | --- |
|  | Mean RT | Hit Rate | d' Sensitivity Index' |
| Difficulty | **2.12 e<sup>+27</sup> \*\*\*** | **3.76 e<sup>+20</sup> \*\*\*** | **2.12 e<sup>+43</sup> \*\*\*** |
| Condition | **0.261 \*** | 0.367 | **0.173\*** |
| Group | 0.606 | 0.619 | 0.445 |
| Difficulty*Condition | **0.107\*** | **0.116\*** | **0.102\*** |
| Difficulty*Group | **0.138\*** | **0.195\*** | 0.972 |
| Condition*Group | 0.349 | **0.283\*** | **0.286\*** |
| Difficulty*Condition*Group | **0.213\*** | **0.165\*** | **0.180\*** |

RT: Reaction Time. Asterisks refer to the level of evidence as expressed in Table 1

#### fMRI results

Since fatigue induction condition (HCL vs LCL) did not statistically influence behavioral performances and subjective states, we focused on the effects of task difficulty on brain activity in the HCL condition only, as our hypothesis was that this condition is the one promoting fatigue induction given the high cognitive load it implies. Nevertheless, we also investigated the HCL vs LCL contrast, in order to detect if fatigue induction condition would modulate brain activity despite its similar effects on behavior.

Brain activity independent from FSMC score was compared between groups (Figure 1B) with F-tests on the following contrasts: HCL vs. LCL, HCL 1-Back, HCL 2-Back, HCL 3-Back, HCL 2-Back vs. HCL 1-Back, HCL 3-Back vs. HCL 1-Back. None of the contrasts showed significant between-group differences in brain activity.

Brain activity correlated to trait fatigue score (FSMC) was assessed for the same six contrasts with F-tests (Figure 1D). Results are displayed in table 5, along with the directionality of the relationship as assessed graphically (see examples on figure S2).

**Table 5.** Results of fMRI analysis: brain regions with activity correlated to the FSMC score.

| Cluster Size<br>(voxels) | *p*<sub>FWE-corrected</sub><br>(Cluster) | Peak *z* | *p*<sub>FWE-corrected</sub><br>(Peak) | X Y Z | Regions | Correlation |
| --- | --- | --- | --- | --- | --- | --- |
| Contrast 1 – HCL vs. LCL |  |  |  |  |  |  |
| **No significant voxel** |  |  |  |  |  |  |
| Contrast 2 – HCL 1-Back |  |  |  |  |  |  |
| 2 | 0.075 | 3.50 | 0.032* | -48 -40 56 | Left intraparietal sulcus | Pos in pwMS |
| 1 | 0.089 | 3.44 | 0.040* | -39 -28 -16 | Left inferior temporal sulcus | Pos in HC |
| Contrast 3 – HCL 2-Back |  |  |  |  |  |  |
| 5 | 0.017* | 3.79 | 0.004** | -48 -13 -13 | Left superior temporal sulcus | Pos in HC<br>Neg in pwMS |
| 7 | 0.014* | 4.07 | 0.001** | -42 -25 -16 | Left inferior temporal sulcus | Pos in HC |
| 16 | 0.007** | 3.87 | 0.003** | -3 -55 23 | Left precuneus/retrosplenial cortex | Pos in HC |
| 9 | 0.012* | 3.62 | 0.007** | -15 -82 -34 | Left and right cerebellum | Pos in HC |
| 5 | 0.017* | 3.24 | 0.024* | 30 -79 -34 |  |  |
| Contrast 4 – HCL 3-Back |  |  |  |  |  |  |
| 17 | 0.009** | 4.86 | <0.001*** | -42 -25 -16 | Left inferior temporal sulcus | Pos in HC |
| 5 | 0.026* | 3.32 | 0.029* | -3 -55 23 | Left precuneus/retrosplenial cortex | Pos in HC |
| 5 | 0.026* | 4.00 | 0.002** | -15 -85 -34 | Left Cerebellum | Pos in HC |
| 1 | 0.045* | 3.53 | 0.014* | -39 -73 -37 |  | Neg in MS |
| Contrast 5 – HCL 2-Back vs. HCL 1-Back |  |  |  |  |  |  |
| 1 | 0.022* | 3.44 | 0.007** | -3 41 56 | Left superior frontal gyrus | Pos in HC<br>Neg in MS |
| 6 | 0.018* | 3.75 | 0.002** | -33 2 29 | Left precentral sulcus | Pos in HC<br>Neg in MS |
| 8 | 0.017* | 3.92 | 0.001** | -48 -13 -13 | Left inferior temporal sulcus | Pos in HC |
| 16 | 0.012* | 3.93 | 0.001** | -6 -31 35 | Left mid cingulate | Pos in HC<br>Neg in MS |
| 1 | 0.022* | 3.36 | 0.010* | 27 -79 -31 | Right cerebellum | Pos in HC |
| Contrast 6 – HCL 3-Back vs. HCL 1-Back |  |  |  |  |  |  |
| 8 | 0.026* | 3.86 | 0.005** | 15 -37 59 | Right postcentral gyrus | Pos in HC<br>Neg in MS |
| 4 | 0.040* | 3.64 | 0.012* | -36 2 26 | Left precentral sulcus | Pos in HC<br>Neg in MS |
| 4 | 0.040* | 3.52 | 0.018* | -48 -16 -13 | Left inferior temporal sulcus | Pos in HC<br>Neg in MS |
| 1 | 0.059 | 3.25 | 0.047* | 9 -10 50 | Cingulate motor area | Pos in HC |
| 12 | 0.019* | 3.56 | 0.017* | -15 -31 8 | Left pulvinar thalamus | Pos in HC<br>Neg in MS |
| 3 | 0.044* | 3.44 | 0.025* | 0 -1 8 | Right anterior thalamus | Pos in HC<br>Neg in MS |
| 4 | 0.040* | 3.30 | 0.040* | 18 -4 23 | Right Caudate | Neg in MS |
| 5 | 0.035* | 3.37 | 0.031* | -36 -76 -34 | Left Cerebellum | Pos in HC<br>Neg in MS |

HCL: High Cognitive Load condition; LCL: Low Cognitive Load Condition; Pos: positive correlation; Neg: negative correlation; pwMS: person with multiple sclerosis; HC: healthy controls. \**p* < 0.05; \*\**p* < 0.01; \*\*\**p* < 0.001

Evaluation of fatigue induction condition (HCL vs. LCL) yielded no significant result. However, analysis of each N-Back difficulty level (figure 3, left panel) showed that the easiest condition (1-Back) led to a positive association between the FSMC score and the left intraparietal sulcus (IPS) in pwMS, and in the left inferior temporal sulcus (ITS) in HC (figure 3A). During the 2-Back condition (figure 3B), brain activity in HC was positively correlated to FSMC in left ITS, left parietal areas (precuneus and retrosplenial cortex), and cerebellum bilaterally. An interaction was found in the left superior temporal sulcus (STS), where brain activity was positively correlated with fatigue score in controls, and negatively in pwMS. For the 3-back condition (figure 3C), brain activity in left ITS and parietal areas was positively correlated to FSMC score in HC and the left cerebellum showed an interaction effect (positive correlation in HC, negative in pwMS).

When taking explicitly into account 1-Back condition as the reference level (figure 3, right panel), we observed for medium difficulty (2-Back > 1-Back) a positive association between FSMC score and brain activity in the left ITS and right cerebellum in HC (figure 3D). Interaction effects (correlation with FSMC score positive in HC and negative in pwMS) were observed in regions belonging to the sensorimotor and associative STC networks (left midcingulate, left precentral sulcus and left superior frontal gyrus). For the highest difficulty level (3-Back > 1-Back), the same interaction effect was observed in the STC network (left precentral, right postcentral sulcus, and thalamus bilaterally), as well as in left ITS and the cerebellum (figure 3E). Two additional regions of the STC network, the right caudate and the right cingulate motor area were also linked to the fatigue score, negatively in the pwMS for the former and positively in HC for the latter.

**Figure 3.**
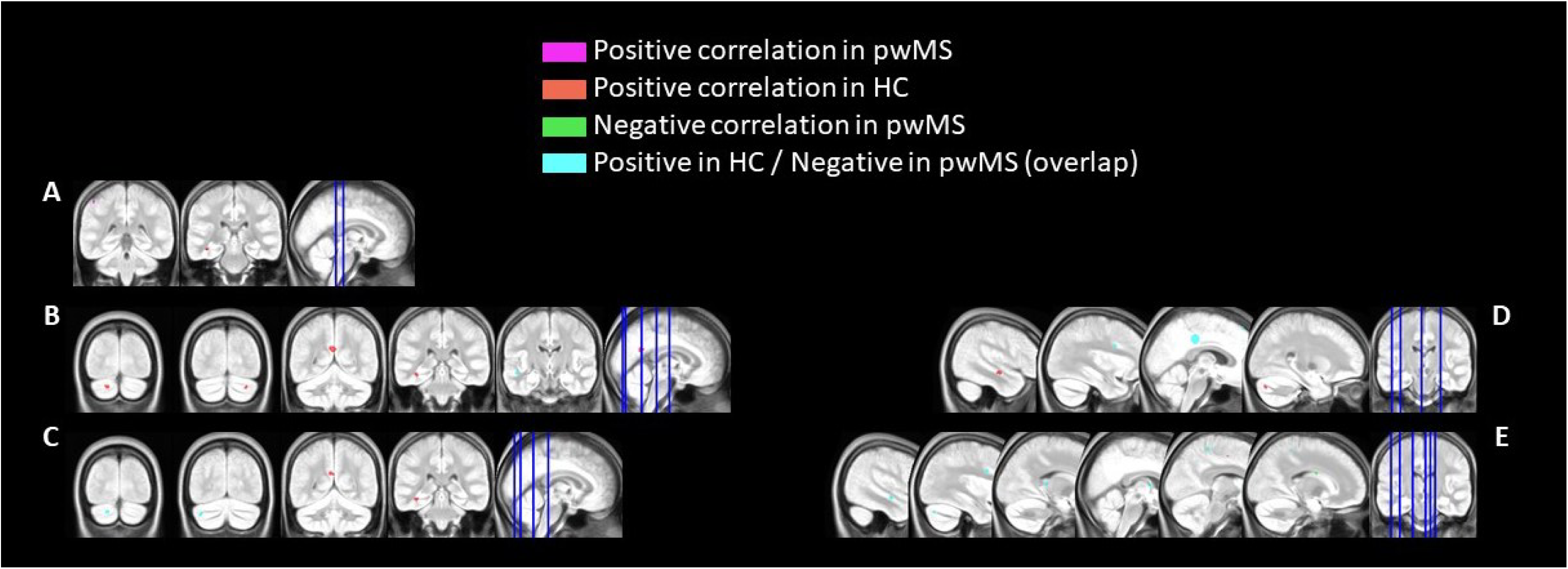
Group-specific clusters of brain regions whose activity correlated to FSMC score during. Clusters of brain regions whose activity is positively correlated to FSMC in pwMS only (purple), HC only (red), negatively in pwMS only (green) and with a between-group interaction (positive correlation in HC and negative in pwMS, cyan) for the 1-Back (A), 2-Back (B) and 3-Back (C) conditions and the 2-Back>1-Back (D) and the 3-Back>1-Back (E) contrasts.

## Discussion

Following a fatigue induction protocol, brain activity was recorded during an N-Back task in pwMS and HC. Both groups showed a similar increase of subjective fatigue following fatigue induction. Between-group differences were observed neither for performance at the N-Back task nor in terms of brain responses independent of fatigue. By contrast, trait fatigue modulated selectively brain responses in one group but not another, mainly in sensorimotor and associative regions of the STC network.

### Behavioral data

Fatigue prevalence in the MS group is in line with the literature (Weiland et al., 2015), with 75% of our pwMS experiencing at least mild fatigue. Yet, there exists no between-group difference regarding trait fatigue, anxiety, depression and sleep quality. As participants were recruited between 2018 and 2021, the deleterious impact of COVID-19 pandemic and lockdowns on mood, fatigue and sleep in the general population could explain scores in the HC group (Cellini et al., 2021; Charonitis et al., submitted.).

As expected, the TLDB task induced a subjective fatigue state until the end of data acquisition, and higher difficulty level at the N-back task is associated with slower response times and lower accuracy. However, we did not observe an effect of fatigue induction condition on performance. Contrary to previous studies, the effect of fatigue induction on working memory was measured as the consequence of the recruitment of more or less extensive cognitive resources, and not by comparison to a passive condition (e.g., rest or watching videos, as for instance in Gilsoul et al., in prep, and Spiteri et al., 2017). Based on subjective scale data, we interpreted the absence of difference between HCL and LCL conditions as reflecting a similar effect of our two conditions on subsequent working memory performance.

Strikingly, we observed similar effects of fatigue induction and task difficulty in the two groups. Like previously reported by Borragán et al. (2018) in mildly impaired patients, our pwMS in the early stage of the disease do not show increased fatigue susceptibility by comparison to HC when the task is adjusted to processing capacity. However, we do not observe more fatigue in the HCL than LCL condition, which we interpret as an effect of task’s length, independent of cognitive load (Guillemin et al., 2022).

Not only did we not observe group effects on N-Back performance, but we provide evidence for an absence of effect for the interaction between difficulty level and group for RTs and Hit Rate. This result suggests that pwMS did not show an increased alteration of performance as difficulty increased compared to HC. Based on this evidence, we consider that pwMS and HC performed the N-back task in a globally similar way. However, as we focused on pwMS in the early stage of the disease, alterations of performance might be subtle.

### Group-Specific Fatigue Substrates According to Difficulty Level

We did not observe differences in brain response between pwMS and HC during performance at the N-Back task, while there exist group-specific associations between fatigue score and brain activity for the three difficulty levels. Positive associations are observed in HC in the left ITS (1B, 2B, 3B) and precuneus (2B, 3B). In our study, the ITS was implicated in fatigued HC in every difficulty level. Inferior temporal regions are usually found in verbal working memory tasks (Buchsbaum & D’Esposito, 2019) and we had no a-priori hypothesis on its association with fatigue in MS. Regarding the precuneus, it is a region belonging to the default mode network that is usually deactivated during a cognitive task, especially as workload increases (Chen et al., 2020). However, this region has also been associated with awareness and self-reflection (Cavanna, 2007), and its activation in a context of fatigue could reflect perception of exertion (Fontes et al., 2015).

During the 1-Back condition, fatigue was positively correlated to activity in the intraparietal sulcus (IPS) in pwMS. The IPS is the core region of the dorsal attention network (DAN), which is principally involved in top-down voluntary orientation of attention (Asplund, Todd, Snyder, & Marois, 2010; Chica, Bartolomeo, & Valero-Cabré, 2011). The 1-Back task is usually associated with a recruitment of the ventral attention network and stimulus driven attention in young healthy subjects (Kurth et al., 2016). Consequently, we propose that the recruitment of the DAN in our early pwMS reflects an additional effort to perform the task, which could be implicated in fatigue genesis.

Activity in the cerebellum was associated to FSMC positively in HC during the 2-and 3-Back conditions, and negatively in pwMS during the most difficult task level. The cerebellum integrates internal and external stimulations to maintain homeostasis and performance through automatization of cognitive processes (Schmahmann, 2019). We suggest that increased cerebellar activity in fatigued HC could reflect an enhanced automatization strategy to preserve performance (Bonnet et al., 2010). In pwMS, the result obtained is in line with other study showing a decreased cerebellar activity during cognitive tasks of high load (Bonnet et al., 2010; Chen et al., 2020), and in fatigued patients (Filippi et al., 2002). Like previously proposed by Bonnet et al. (2010), we suggest that a decreased implication of the cerebellum relates to a difficulty in pwMS to generate automated attentional procedure, which could in turn trigger fatigue symptom (for example due to the necessity to higher recruitment of the DAN network to maintain performance).

### Implication of the Striato-Thalamo-Cortical Network in Early MS Fatigue

To better delineate the effects of fatigue on working memory, we compared brain activity in the 2 and 3-Back blocks to 1-Back blocks (involving mainly perceptual and motor requirements). Correlations (positive in HC and negative in pwMS) in the cerebellum and the inferior temporal sulcus remained significant, but regions belonging to the STC network also emerged: the thalami, the cingulate cortex, the basal ganglia (right caudate) and the fronto-parietal cortex, providing additional evidence for Chaudhuri & Behan’s model of fatigue (Chaudhuri & Behan, 2000). In line with their model, we propose that in fatigued HC, the increased recruitment of regions from the STC reflects a compensatory mechanism to sustain motivation and maintain performance, while in pwMS, the disruption of the several STC loops (at least sensorimotor and associative as evidenced in this study) will generate fatigue. Interestingly, the fatigue-related activity is modulated by task-difficulty, with the involvement of basal ganglia and the thalamus in the 3B vs. 1B contrast, while only cortical regions from the STC network were implicated in the 2B vs. 1B. Therefore, we suggest that in early MS, activity of the STC loops might be fatigue-dependent for cognitive tasks of high demand, and becomes implicated in fatigue processes in simpler tasks as disease progresses, as it is the case in other studies (see for example DeLuca, Genova, Hillary, & Wylie, 2008; Rocca et al., 2009).

### Fatigue in early pwMS is linked to widespread functional reorganization

Apart from the easiest task condition (1-Back), where fatigue was positively associated with activity in the IPS in pwMS, brain activity in pwMS was systematically negatively associated with fatigue and occurred in regions where a positive association was observed in HC. Yet, behavioral performances did not differ between groups. MS fatigue has been linked to performance decrement and hyperactivity of numerous brain regions (see ARM et al., 2019 for a review), but none of these studies focused on the beginning of the disease. One study, though, reported that hyperactivity in frontal areas during a motor task in fatigued patients is higher in pwMS with a disease duration over 5 years (Rocca et al., 2016).

It seems thus that in the earliest stages of the disease, fatigue is not necessarily associated with a higher recruitment of brain areas linked to fatigue in HC. We tentatively propose that, at disease onset, structural alterations of the brain connectome will gradually lead to the emergence of alternative functional networks through neuronal plasticity and compensation in order to maintain cognitive performance (Capone, Collorone, Cortese, Di Lazzaro, & Moccia, 2020; Schoonheim, Meijer, & Geurts, 2015). This explains why we observe negative association between fatigue score and recruitment of brain areas previously associated with fatigue. Later on, networks reorganization may stabilize but not be sufficient to prevent cognitive decline, which could explain the results obtained by the abovementioned studies investigating the cerebral correlates of fatigue in pwMS at more advanced stages. This hypothesis is in line with recent longitudinal evidence in CIS patients, showing structural and functional connectivity reorganization throughout indirect anatomical connection with preserved cognitive performance at one year following disease onset (Koubiyr et al., 2019). These early changes were followed by a more constrained reorganization associated with cognitive decline at 5 years from disease onset (Koubiyr et al., 2021).

## Limitations

Several limitations have to be pointed out in the current study. First, the low cognitive load condition also triggered fatigue in our protocol. Therefore, it cannot be considered as a control condition. Our protocol remains relevant to demonstrate that a continuous task of low time constraint can also elicit subjective state of fatigue relatively fast, as most fatigue inducing protocols in the literature lasts much longer (32 mins in our experiment vs. > 1 H in most studies, as for instance in Lorist, 2008; van der Linden, Frese, & Meijman, 2003; Wang, Ding, & Kluger, 2014). However, we were not able to study the effects of trait fatigue on cerebral activity when participants felt rested, or the effect of state fatigue by comparison to non-fatigued participants. Further task fMRI protocols are needed to better investigate if trait and state fatigue are linked to distinct brain substrates in early MS.

Second, it is important to note that our fatigue induction protocol also had a significant impact on sleepiness and motivation, in both groups. Fatigue, sleepiness and motivation are three highly related notions that are sometimes referred to interchangeably in the literature. Yet, they are very distinct concepts. In their study, Borragán et al. (2017), succeeded in inducing fatigue without increasing sleepiness using the original version of the TLDB. Motivation scores were, however, not reported. In fact, most imaging studies in the field of fatigue in MS do not report states of motivation nor sleepiness. Nevertheless, both states might modulate the link observed between fatigue and cerebral activity. Additionally, we did not statistically control for confounding factors such as depression, anxiety and sleep disorder. However, no between group differences were evidenced regarding those secondary causes of fatigue.

As we were interested in studying disease specific brain correlates of fatigue, we did not analyze correlations that are similar in both groups. Such analysis would have explored the common mechanisms of fatigue in healthy subjects and patients, which is also of great interest but was not the aim of the present study. Besides, the sample included in this study was small, and additional disease-specific correlations with smaller effect sizes might not have been evidenced here. Consequently, while the present study provides initial results on the functional cerebral substrates of fatigue in the early stages of the disease, further works should complement this first exploration.

## Conclusion

To conclude, we identified specific activation patterns relating to fatigue in pwMS and HC, despite similar behavioral performance and subjective fatigue. We found additional evidence for the implication of the striato-thalamo-cortical circuitry in fatigue pathophysiology, and that this implication might be task-dependent (i.e. more pronounced in challenging cognitive tasks). Moreover, our results suggest that pwMS in the early stages of the disease recruit an alternative attentional circuit compared to HC to perform the task. We hypothesized that an alteration of automated strategies led to a reorganized pattern of activation in pwMS and that this cerebral reorganization could trigger fatigue. Hence, this is the first task fMRI study indicating an alteration of functional activity in regions of the STC network from the beginning of the disease.

## List of Abbreviations

CIS: Clinically Isolated Syndrome
HC: Healthy Controls
DAN: Dorsal Attention Network
HCL: High Cognitive Load
IPS: Intraparietal Sulcus
ITS: Inferior Temporal Sulcus
LCL: Low Cognitive Load
MS: Multiple Sclerosis
pwMS: people with Multiple Sclerosis
RR: Relapsing-Remitting
STC: Striato-Thalamo-Cortical
STD: Stimulus Time Duration
STS: Superior Temporal Sulcus
TLDB: Time Load Dual Back;

## Acknowledgements

This work was conducted at the GIGA In Vivo Imaging platform of University of Liège, Belgium. We want to thank all the participants who took part of the present study. This work was supported by Fonds National de la Recherche Scientifique (FNRS), University of Liège and the Fauconnier and Sallets Fund from the King Baudouin Fundation (KBF-FRB). C.G. was supported by University of Liège. C.P., F.C. were supported by Fonds National de la Recherche Scientifique (FRS-FNRS), Belgium.

## Statements & Declarations

### Fundings

This work was supported by Fonds National de la Recherche Scientifique (FNRS), University of Liège and the Fauconnier and Sallets Fund from the King Baudouin Fundation (KBF-FRB). C.G. was supported by University of Liège. C.P., F.C. were supported by Fonds National de la Recherche Scientifique (FRS-FNRS), Belgium. None of funding sources had an involvement in study design; in the collection, analysis, interpretation of data; in the writing of the report; in the decision to submit the article for publication.

### Competing interests

The authors have no relevant financial or non-financial interests to disclose.

### Author contribution

<u>Conceptualisation</u>: Camille Guillemin, Emilie Lommers, Pierre Maquet, Fabienne Collette, <u>Methodology</u> : Camille Guillemin, Pierre Maquet, Fabienne Collette ; <u>Investigation</u> : Camille Guillemin, Maëlle Charonitis, Florence Requier, Gaël Delrue ; <u>Software</u> : Camille Guillemin, Nikita Beliy, Christophe Phillips ; <u>Data curation</u> : Camille Guillemin, Maëlle Charonitis, Nikita Beliy ; <u>Formal analysis</u> : Camille Guillemin, Maëlle Charonitis, Christophe Phillips, Fabienne Collette ; <u>Resources</u> : Emilie Lommers ; <u>Funding acquisition</u> : Pierre Maquet, Fabienne Collette ; <u>Project administration</u> : Camille Guillemin ; <u>Supervision</u> : Christophe Phillips, Fabienne Collette ; <u>Validation</u> : Fabienne Collette. The first draft of the manuscript was written by Camille Guillemin and all authors commented on previous versions of the manuscript. All authors read and approved the final manuscript.

### Data availability

The data underlying this report are made available from the corresponding author on request following a formal data sharing agreement.

## Notes

### Competing Interest Statement

The authors have declared no competing interest.

## References

Åkerstedt, T., & Gillberg, M. (1990). Subjective and Objective Sleepiness in the Active Individual. International Journal of Neuroscience, 52(1–2), 29–37. 10.3109/00207459008994241

Alexander, G. E., DeLong, M. R., & Strick, P. L. (1986). Parallel Organization of Functionally Segregated Circuits Linking Basal Ganglia and Cortex. Annual Review of Neuroscience, 9(1), 357–381. 10.1146/annurev.ne.09.030186.002041

Arm, J., Ribbons, K., Lechner-Scott, J., & Ramadan, S. (2019). Evaluation of MS related central fatigue using MR neuroimaging methods: Scoping review. Journal of the Neurological Sciences, 400(August 2018), 52–71. 10.1016/j.jns.2019.03.007

Asplund, C. L., Todd, J. J., Snyder, A. P., & Marois, R. (2010). A central role for the lateral prefrontal cortex in goal-directed and stimulus-driven attention. Nature Neuroscience, 13(4), 507–512. 10.1038/nn.2509

Bakshi, R., Shaikh, Z. A., Miletich, R. S., Czarnecki, D., Dmochowski, J., Henschel, K., & Janardhan, V. (2000). Fatigue in multiple sclerosis and its relationship to depression and neurologic disability. Multiple Sclerosis, 6(February), 181–185.

Beck, A. T., Ward, C. H., Mendelson, M., Mock, J., & Erbaugh, J. (1961). An Inventory for Measuring Depression. Archives of General Psychiatry, 4(6), 561. Retrieved from http://archpsyc.jamanetwork.com/article.aspx?doi=10.1001/archpsyc.1961.01710120031004

Bonnet, M. C., Allard, M., Dilharreguy, B., Deloire, M., Petry, K. G., & Brochet, B. (2010). Cognitive compensation failure in multiple sclerosis. Neurology, 75(14), 1241–1248. 10.1212/WNL.0b013e3181f612e3

Borragán, G., Slama, H., Bartolomei, M., & Peigneux, P. (2017). Cognitive fatigue: A Time-based Resource-sharing account. Cortex, 89, 71–84. 10.1016/j.cortex.2017.01.023

Buchsbaum, B. R., & D’Esposito, M. (2019, March 1). A sensorimotor view of verbal working memory. Cortex. Masson SpA. 10.1016/j.cortex.2018.11.010

Buysse, D. J., Reynolds, C. F., Monk, T. H., Berman, S. R., & Kupfer, D. J. (1998). The Pittsburg Sleep Quality Index: A New Instrument for Psychiatric Practice and Research. Psychiatry Research.

Capone, F., Collorone, S., Cortese, R., Di Lazzaro, V., & Moccia, M. (2020). Fatigue in multiple sclerosis: The role of thalamus. Multiple Sclerosis Journal, 26(1), 6–16. 10.1177/1352458519851247

Cavanna, A. E. (2007). The precuneus and consciousness. CNS Spectrums. MBL Communications. 10.1017/S1092852900021295

Cellini, N., Conte, F., De Rosa, O., Giganti, F., Malloggi, S., Reyt, M., … Ficca, G. (2021). Changes in sleep timing and subjective sleep quality during the COVID-19 lockdown in Italy and Belgium: age, gender and working status as modulating factors. Sleep Medicine, 77, 112–119. 10.1016/j.sleep.2020.11.027

Charonitis, M., Requier, F., Guillemin, C., Reyt, M., Folville, A., Geurten, M., … Collette, F. (submitted). Impact of COVID19 pandemic on fatigue during lockdown and one-year later. En preparation.

Chaudhuri, A., & Behan, P. O. (2000). Fatigue and basal ganglia. Journal of the Neurological Sciences, 179, 34–42. Retrieved from http://www.sciencedirect.com/science/article/pii/S0022510X00004111%0Apapers2://publication/uuid/4E2A9FEC-EEDA-4392-AE1B-98B06EE34D7F

Chaudhuri, A., & Behan, P. O. (2004, October 1). Multiple sclerosis is not an autoimmune disease. Archives of Neurology. American Medical Association. 10.1001/archneur.61.10.1610

Chen, M. H., Wylie, G. R., Sandroff, B. M., Dacosta-Aguayo, R., DeLuca, J., & Genova, H. M. (2020). Neural mechanisms underlying state mental fatigue in multiple sclerosis: a pilot study. Journal of Neurology, 1–11. 10.1007/s00415-020-09853-w

Chiaravalloti, N. D., & DeLuca, J. (2008). Cognitive impairment in multiple sclerosis. The Lancet Neurology, 7(12), 1139–1151. 10.1016/S1474-4422(08)70259-X

Chica, A. B., Bartolomeo, P., & Valero-Cabré, A. (2011). Dorsal and ventral parietal contributions to spatial orienting in the human brain. Journal of Neuroscience, 31(22), 8143–8149. 10.1523/JNEUROSCI.5463-10.2010

DeLong, M., & Wichmann, T. (2010). Changing views of basal ganglia circuits and circuit disorders. Clinical EEG and Neuroscience, 41(2), 61–67. 10.1177/155005941004100204

DeLuca, J., Genova, H. M., Hillary, F. G., & Wylie, G. (2008). Neural correlates of cognitive fatigue in multiple sclerosis using functional MRI. Journal of the Neurological Sciences, 270(1–2), 28–39. 10.1016/j.jns.2008.01.018

Filippi, M., Rocca, M. A., Colombo, B., Falini, A., Codella, M., Scotti, G., & Comi, G. (2002). Functional Magnetic Resonance Imaging Correlates of Fatigue in Multiple Sclerosis. NeuroImage, 15(3), 559–567. 10.1006/nimg.2001.1011

Fontes, E. B., Okano, A. H., De Guio, F., Schabort, E. J., Min, L. L., Basset, F. A., … Noakes, T. D. (2015). Brain activity and perceived exertion during cycling exercise: An fMRI study. British Journal of Sports Medicine, 49(8), 556–560. 10.1136/bjsports-2012-091924

Gilsoul, J., Charonitis, M., Bahri, M. A., Phillips, C., Depierreux, F., Salmon, E., … Collette, F. (in preparation). Age-related changes in cerebral activity following cognitive fatigue during a working memory task: an fMRI investigation.

Gorgolewski, K. J., Auer, T., Calhoun, V. D., Craddock, R. C., Das, S., Duff, E. P., … Poldrack, R. A. (2016). The brain imaging data structure, a format for organizing and describing outputs of neuroimaging experiments. Scientific Data, 3. 10.1038/sdata.2016.44

Guillemin, C., Hammad, G., Read, J., Requier, F., Charonitis, M., Delrue, G., … Collette, F. (2022). Pupil response speed as a marker of cognitive fatigue in early Multiple Sclerosis☆. Multiple Sclerosis and Related Disorders, 65(May), 104001. 10.1016/j.msard.2022.104001

Jamil, T., Ly, A., Morey, R. D., Love, J., Marsman, M., & Wagenmakers, E. J. (2017). Default “Gunel and Dickey” Bayes factors for contingency tables. Behavior Research Methods, 49(2), 638–652. 10.3758/s13428-016-0739-8

Jeffreys, H. (1961). Theory of Probability. 3rd Edition. Oxford: Clarendon Press.

Johns, M. W. (1991). A New Method for Measuring Daytime Sleepiness: The Epworth Sleepiness Scale. Sleep, 14(6), 540–545. 10.1093/sleep/14.6.540

Keysers, C., Gazzola, V., & Wagenmakers, E.-J. (2020). Using Bayes factor hypothesis testing in neuroscience to establish evidence of absence. Nature Neuroscience, 23(7), 788–799. 10.1038/s41593-020-0660-4

Kirchner, W. K. (1958). Age differences in short-term retention of rapidly changing information. Journal of Experimental Psychology, 55(4), 352–358. 10.1037/h0043688

Koubiyr, I., Besson, P., Deloire, M., Charre-Morin, J., Saubusse, A., Tourdias, T., … Ruet, A. (2019). Dynamic modular-level alterations of structural-functional coupling in clinically isolated syndrome. Brain, 142(11), 3428–3439. 10.1093/brain/awz270

Koubiyr, I., Deloire, M., Brochet, B., Besson, P., Charré-Morin, J., Saubusse, A., … Ruet, A. (2021). Structural constraints of functional connectivity drive cognitive impairment in the early stages of multiple sclerosis. Multiple Sclerosis Journal, 27(4), 559–567. 10.1177/1352458520971807

Krupp, L. B., Alvarez, L. A., LaRocca, N. G., & Scheinberg, L. C. (1988). Fatigue in Multiple Sclerosis. Arch. Neurol., 45, 435–437. Retrieved from http://archneur.jamanetwork.com/

Kurth, S., Majerus, S., Bastin, C., Collette, F., Jaspar, M., Bahri, M. A., & Salmon, E. (2016). Effects of aging on task-and stimulus-related cerebral attention networks. Neurobiology of Aging, 44, 85–95. 10.1016/j.neurobiolaging.2016.04.015

Kurtzke, J. F. (1983). Rating neurologic impairment in multiple sclerosis: An expanded disability status scale (EDSS). Neurology, 33(11), 1444–1452. 10.1212/wnl.33.11.1444

Lee, K. A., Hicks, G., & Nino-Murcia, G. (1991). Validity and reliability of a scale to assess fatigue. Psychiatry Research, 36(3), 291–298. 10.1016/0165-1781(91)90027-M

Lorist, M. M. (2008). Impact of top-down control during mental fatigue. Brain Research, 1232, 113–123. 10.1016/j.brainres.2008.07.053

Newland, P., Starkweather, A., & Sorenson, M. (2016). Central fatigue in multiple sclerosis: a review of the literature. Journal of Spinal Cord Medicine. Taylor & Francis. 10.1080/10790268.2016.1168587

Palotai, M., & Guttmann, C. R. G. (2020, June 1). Brain anatomical correlates of fatigue in multiple sclerosis. Multiple Sclerosis Journal. SAGE Publications Ltd. 10.1177/1352458519876032

Penner, I. K., Raselli, C., Stöcklin, M., Opwis, K., Kappos, L., & Calabrese, P. (2009). The Fatigue Scale for Motor and Cognitive Functions (FSMC): Validation of a new instrument to assess multiple sclerosis-related fatigue. Multiple Sclerosis, 15(12), 1509–1517. 10.1177/1352458509348519

Peters, S. K., Dunlop, K., & Downar, J. (2016, December 27). Cortico-striatal-thalamic loop circuits of the salience network: A central pathway in psychiatric disease and treatment. Frontiers in Systems Neuroscience. Frontiers Media S.A. 10.3389/fnsys.2016.00104

Phillips, C. U. de L.-Ul. > G.-C. I. vivo I., & Pernet, C. U. of E. (2018). *Unifying lesion masking and tissue probability maps for improved segmentation and normalization*. *23rd Annual Meeting of the Organization for Human Brain Mapping*. Retrieved from https://github.com/CyclotronResearchCentre/USwithLesion.

Posner, J., Marsh, R., Maia, T. V., Peterson, B. S., Gruber, A., & Simpson, H. B. (2014). Reduced functional connectivity within the limbic cortico-striato-thalamo-cortical loop in unmedicated adults with obsessive-compulsive disorder. Human Brain Mapping, 35(6), 2852–2860. 10.1002/hbm.22371

Rǎdulescu, A., Herron, J., Kennedy, C., & Scimemi, A. (2017). Global and local excitation and inhibition shape the dynamics of the cortico-striatal-thalamo-cortical pathway. Scientific Reports, 7(1), 1–21. 10.1038/s41598-017-07527-8

Rocca, M. A., Gatti, R., Agosta, F., Broglia, P., Rossi, P., Riboldi, E., … Filippi, M. (2009). Influence of task complexity during coordinated hand and foot movements in MS patients with and without fatigue : AA kinematic and functional MRI study. Journal of Neurology, 256(3), 470–482. 10.1007/s00415-009-0116-y

Rocca, Maria A., Meani, A., Riccitelli, G. C., Colombo, B., Rodegher, M., Falini, A., … Filippi, M. (2016). Abnormal adaptation over time of motor network recruitment in multiple sclerosis patients with fatigue. Multiple Sclerosis, 22(9), 1144–1153. 10.1177/1352458515614407

Roelcke, U., Kappos, L., Lechner-Scott, J., Brunnschweiler, H., Huber, S., Ammann, W., … Leenders, K. L. (1997). Reduced glucose metabolism in the frontal cortex and basal ganglia of multiple sclerosis patients with fatigue: A 18F-fluorodeoxyglucose positron emission tomography study. Neurology, 48(6), 1566–1571. 10.1212/WNL.48.6.1566

Runia, T. F., Jafari, N., Siepman, D. A. M., & Hintzen, R. Q. (2015). Fatigue at time of CIS is an independent predictor of a subsequent diagnosis of multiple sclerosis. *Journal of Neurology*, Neurosurgery and Psychiatry, 86(5), 543–546. 10.1136/jnnp-2014-308374

Schmahmann, J. D. (2019). The cerebellum and cognition. Neuroscience Letters, 688(July 2018), 62–75. 10.1016/j.neulet.2018.07.005

Schmidt, P., Gaser, C., Arsic, M., Buck, D., Förschler, A., Berthele, A., … Mühlau, M. (2012). An automated tool for detection of FLAIR-hyperintense white-matter lesions in Multiple Sclerosis. NeuroImage, 59(4), 3774–3783. 10.1016/j.neuroimage.2011.11.032

Schoonheim, M. M., Meijer, K. A., & Geurts, J. J. G. (2015). Network Collapse and Cognitive Impairment in Multiple Sclerosis. Frontiers in Neurology, 6, 82. 10.3389/fneur.2015.00082

Spielberger, C. D., Gorsuch, R. L., Ushene, R., Vagg, P. R., & Jacobs, G. A. (1983). Manual for the State-Trait Anxiety Inventory. Palo Alto, CA: Consulting Psychologists Press.

Spiteri, S., Hassa, T., Claros-Salinas, D., Dettmers, C., & Schoenfeld, M. A. (2017). Neural correlates of effort-dependent and effort-independent cognitive fatigue components in patients with multiple sclerosis. Multiple Sclerosis Journal, 1–11. 10.1177/1352458517743090

Thompson, A. J., Banwell, B. L., Barkhof, F., Carroll, W. M., Coetzee, T., Comi, G., … Cohen, J. A. (2018). Diagnosis of multiple sclerosis: 2017 revisions of the McDonald criteria. The Lancet Neurology, 17(2), 162–173. 10.1016/S1474-4422(17)30470-2

Tzourio-Mazoyer, N., Landeau, B., Papathanassiou, D., Crivello, F., Etard, O., Delcroix, N., … Joliot, M. (2002). Automated anatomical labeling of activations in SPM using a macroscopic anatomical parcellation of the MNI MRI single-subject brain. NeuroImage, 15(1), 273–289. 10.1006/nimg.2001.0978

van der Linden, D., Frese, M., & Meijman, T. F. (2003). Mental fatigue and the control of cognitive processes: Effects on perseveration and planning. Acta Psychologica. 10.1016/S0001-6918(02)00150-6

Wang, C., Ding, M., & Kluger, B. M. (2014). Change in intraindividual variability over time as a key metric for defining performance-based cognitive fatigability. Brain and Cognition, 85(1), 251–258. 10.1016/j.bandc.2014.01.004

Weiland, T. J., Jelinek, G. A., Marck, C. H., Hadgkiss, E. J., van der Meer, D. M., Pereira, N. G., & Taylor, K. L. (2015). Clinically Significant Fatigue: Prevalence and Associated Factors in an International Sample of Adults with Multiple Sclerosis Recruited via the Internet. PLoS ONE, 10(2). 10.1371/journal.pone.0115541

Westfall, P., Johnson, W., & Utts, J. (1997). A Bayesian Perspective on the Bonferroni Adjustment. Biometrika, 84(2).

