## Supplemental Figures for "The specificity of fatigue cerebral substrates in early Multiple Sclerosis - An fMRI study"

### Supplemental Material

**Figure S1. Block-design fMRI N-Back Task**

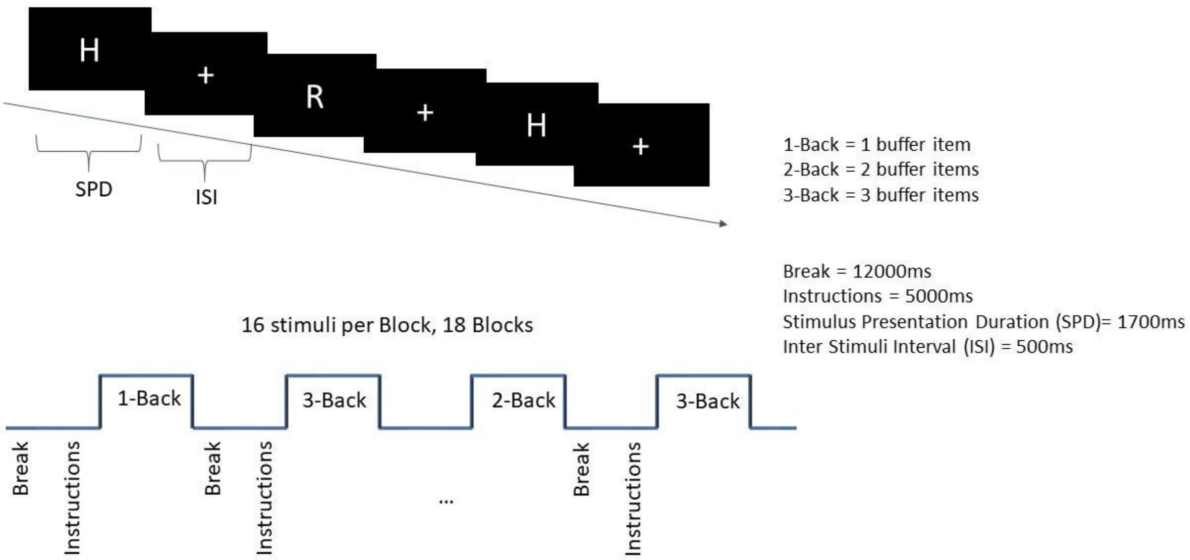


During the N-Back task, each block started with block difficulty (1-Back, 2-Back or 3-Back) instructions displayed during 5000ms on the screen regarding the upcoming block difficulty and a reminder of the corresponding response-keys. In each block, letters were presented with a time presentation (SPD) of 1700ms followed by an inter-stimulus fixation cross (ISI) of 500ms. At the end of the block, a fixation cross of 12000ms (break) was displayed before the beginning of the next block. The first one-to-three letters were considered as "buffer" trials, during which no answer was expected (1 buffer at the beginning of "1-Back" blocks, 2 for the "2-Back", and 3 for the "3-Back"). The total task duration was of 16 minutes, approximately.

**Figure S2. Visual investigation of correlations directionality**

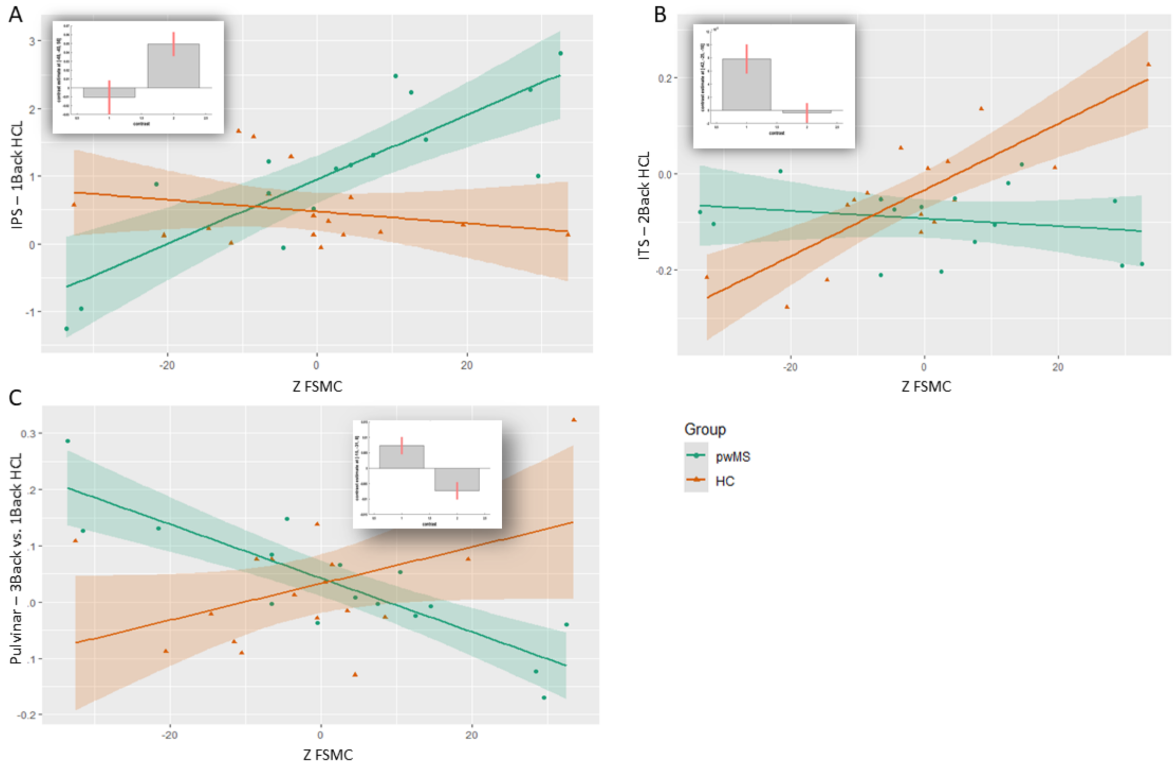


Examples of investigation of correlation directionality. Bar plots represent confidence intervals, left bar plots for healthy controls (HC) and right bar plots for people with Multiple Sclerosis (pwMS). Data points and regression lines are displayed for HC (orange) and pwMS (green) between activity in a peak voxel and mean-centered trait fatigue score (FMSC). A: positive correlation in pwMS in the IPS during 1-Back; B: positive correlation in HC in the ITS during 2-Back; C: positive correlation in HC and negative in pwMS in the Pulvinar during 3-Back vs. 1-Back.

**Figure S3. Evolution of subjective state (fatigue, sleepiness and motivation)**

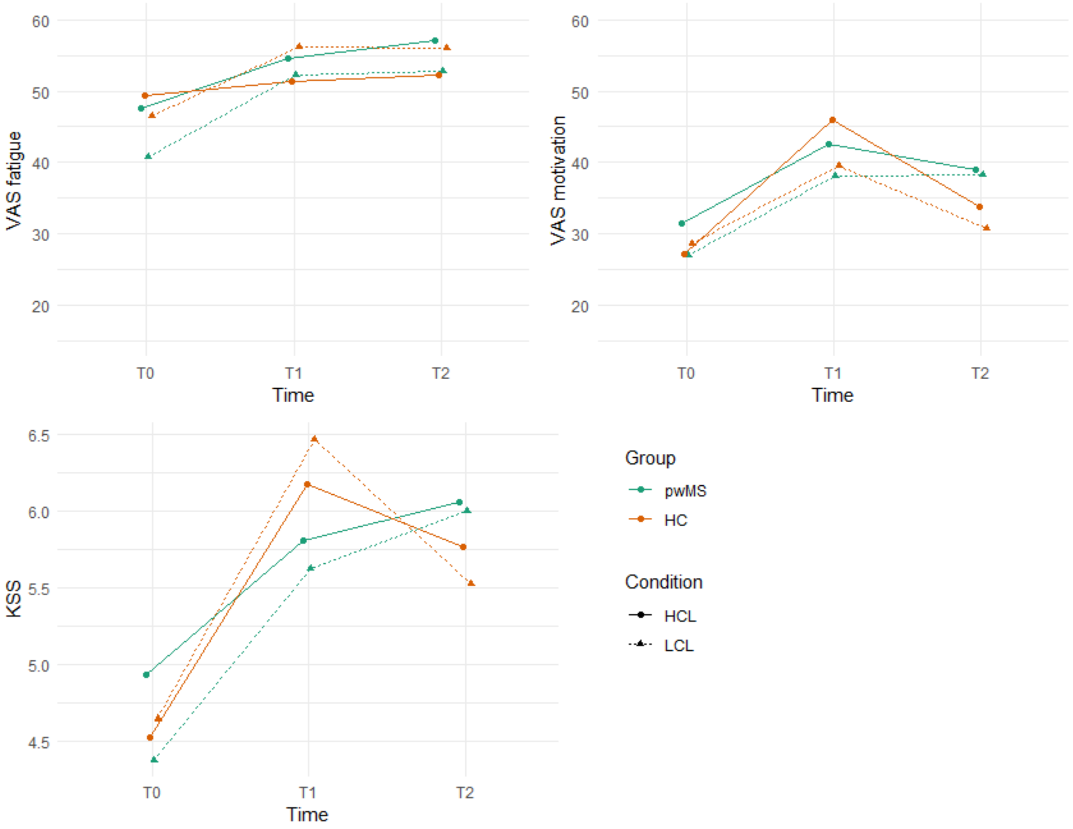


Score obtained for the subjective scales (fatigue and motivation, upper panel, sleepiness, lower panel) before fatigue induction, following fatigue induction and following the N-Back task (T0 to T2). For VAS motivation, a high score corresponds to low motivation. Patients are depicted in green, HC in orange. Solid lines: high cognitive load fatigue induction; Dash lines: low cognitive load fatigue induction.
